# Segregation, Finite Time Elastic Singularities and Coarsening in Renewable Active Matter

**DOI:** 10.1101/2024.09.05.611571

**Authors:** Ayan Roychowdhury, Saptarshi Dasgupta, Madan Rao

**Author notes:** Joint first author (Ayan Roychowdhury, Saptarshi Dasgupta).

## Abstract

Material renewability in active living systems, such as in cells and tissues, can drive the large-scale patterning of forces, with distinctive phenotypic consequences. This is especially significant in the cell cytoskeleton, where multiple species of myosin bound to actin, apply differential contractile stresses and undergo differential turnover, giving rise to patterned force channeling. Here we study the dynamical patterning of stresses that emerge in a hydrodynamic description of a renewable active actomyosin elastomer comprising two myosin species. Our analytical framework also holds for an actomyosin elastomer with a single myosin species. We find that a uniform active contractile elastomer spontaneously segregates into spinodal stress patterns, followed by a finite-time collapse into tension carrying singular structures that display self-similar scaling and caustics. Our numerical analysis carried out in 1D, shows that these singular structures move and merge, and gradually result in a slow coarsening dynamics. We discuss the implications of our findings to the emergence of stress fibers and the spatial patterning of actomyosin. Our study suggests, that with state-dependent turnover of crosslinkers and myosin, the *in vivo* cytoskeleton can navigate through the space of material parameters to achieve a variety of functional phenotypes.

## I. INTRODUCTION

Since actomyosin is the primary agency of cell and tissue scale forces, its patterning along system spanning stress fibers [1], cables or arcs [2, 3] reveals the patterning of forces along tension chains. The cytoskeleton represent a class of active matter called *renewable active matter*; renewability is a distinctive feature of living materials that allows it to navigate through the space of material parameters, resulting in unusual mechanical responses [4] and system spanning force channeling [5]. Here we study the nature of dynamical force patterning using a hydrodynamic description of a renewable active material.

The actomyosin cytoskeleton is put together by the nonequilibrium self-assembly of a variety of crosslinkers and myosin species, that both exert and sense forces, the latter via their strain dependent turnover [6–12]. We will refer to these units of mechanotransduction as *stresslets*; the spatiotemporal patterning of these stresslets mark the patterning of forces. High resolution microscopy reveals that distinct actomyosin stress fibers [2, 3, 5, 13–15], are associated with different stresslet species displaying different cellular localisations [15, 16].

Indeed, the human genome encodes around 40 myosin genes, grouped into 18 classes, with class II being the most prominent in muscle and non-muscle cells [17, 18]. Non-muscle myosin II (NM II), found in all non-muscle eukaryotic cells, assembles into bipolar filaments of up to 30 hexamers [13, 16, 19]. Mammalian cells express three NM II isoforms—NM IIA, NM IIB, and NM IIC— that share identical light chains but differ in their heavy chains, conferring distinct mechanical properties [15, 19–21]. These myosin isoforms differ in their contractile action [22, 23], duty ratio [24] and binding-unbinding rates [15, 19]. Their coordinated interplay is crucial for cellular tension generation and maintenance [15, 16], with the spatially regulated relative abundance leading to strong subcellular localization. In particular, NM IIA and NM IIB are known to undergo self-sorting [15, 25], leading to spatially organized and functionally distinct contractile zones within the cell.

Several studies have explored pattern formation in actomyosin systems, treating them either as active fluids [25–34] or as active elastomers with a single myosin species [35–37]. In contrast to the aster-like patterns and oscillations typical of active fluids, the long-range marginalized elastic interactions in active solids can generate system-spanning stress singularities [4]. The present work goes beyond a routine extension of these studies to multi-species myosin systems; it focuses on how differential stresslet activity and strain dependent turnover can efficiently remodel the effective elastic energy landscape—offering a new way of looking at evolving materials navigating the space of material moduli and driving towards and maintaining itself in mechanically adaptable states [38].

In this paper, we explore the unusual features of such renewable active matter, that exhibits dynamic material renormalization, leading to segregation and singular stress patterns. We construct active hydrodynamic equations [39] for a mixture of contractile stresslets, such as myosin IIA and IIB [13, 15, 20, 21], on a permanently crosslinked actin elastomer, while allowing turnover of myosin stresslets [35–37]. Since the action (stress generation) and reaction (stress sensing dependent turnover) are regulated by independent biochemical cycles, the underlying hydrodynamics is nonreciprocal [40]. A straight-forward material consequence of this nonreciprocity is a renormalization of the elastic moduli that can in principle drive the material to elastic marginality [4]. We find that a slight difference between the contractile activities or turnover rates of the stresslets, can drive spontaneous segregation, *even in the absence of any attractive interaction* – the myosin species with higher contractile activity tend to cluster together (a similar analysis for a single myosin species, leads to a segregation into myosin-rich and myosin-poor regions [41]). However, unlike conventional segregation driven by gradients in chemical potential [42], the spontaneous segregation instability of the myosin stresslets is driven by an effective elastic stress relaxation. At later times, the growing segregated domains collapse into well separated, singular structures of enhanced contractility (tensile “stress fibers”), in striking departure from conventional coarsening. We derive scaling forms for these finite time singular structures [43] and verify them with careful numerics in 1D. The amplification of contractile stresses in the singular tension structures recalls the study in [44]. These singular tension structures can be static or moving [5] and we determine the conditions for these. The moving singular tension structures merge over time and exhibit a slow coarsening dynamics at late times controlled by the differential myosin turnover.

Recent experiments done on cells plated on micro-patterned substrates [5] provide the ideal setup to test our predictions and explore new possibilities on the disassembly of stress fibers upon treatment with the Rho kinase inhibitor Y27632, and their reassembly dynamics following washout of the inhibitor [5, 13, 22]. This provides a perfect experimental platform to carry out systematic quantitative studies on how myosin activation can drive the segregation of an initially homogeneous actomyosin network into domains with differential contractility, and how the highly contractile domains subsequently collapse to form cell-spanning actomyosin cables or stress fibers. In the present setting of continuum elasticity, such taut fibers or cables manifest as singularities or localized concentrations in the tensile stress field along lines (or as “force punctae” in one dimension, as illustrated in this work).

Details of the derivation of the hydrodynamic equations, linear stability analysis and dispersion curves, finite time scaling and convergence analysis, and Movie captions are presented in the Appendix.

## II. HYDRODYNAMIC DESCRIPTION OF ACTOMYOSIN CYTOSKELETON

We work with a 2-dimensional permanently crosslinked actin elastomer of mass density *ρ*_*a*_, embedded in the cytosol, described by a linearized strain tensor ***ϵ*** := (∇***u***+ ∇***u***^*T*^)*/*2, where ***u*** represents the displacement relative to an unstrained reference state. The hydrodynamic linear momentum balance of the overdamped elastomer is 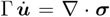, where Γ is the friction of the elastomer with respect to the fluidic cytosol (whose dynamics we neglect since the volume fraction of the mesh is high), and ***σ***(***ϵ***, *ρ*_*a*_, {*ρ*_*i*_}) is the total stress in the elastomer, which depends on the strain ***ϵ***, the density of the actin mesh *ρ*_*a*_, and the densities *ρ*_*i*_ of the *i*^th^ species of active bound myosin (stresslets).

The total stress ***σ*** is the summation of elastic stress ***σ***^*e*^, the viscous stress ***σ***^*d*^, and active stress ***σ***^*a*^: ***σ*** = ***σ***^*e*^+***σ***^*d*^+ ***σ***^*a*^ [35–37]. The elastic stress 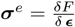 is computed from a free-energy functional 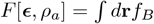, describing an isotropic elastomer, 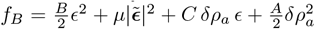, where *ϵ* := tr***ϵ*** is the dilational strain and 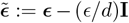 is the deviatoric strain, *B >* 0 and *µ >* 0 are the passive elastic bulk and shear moduli, respectively,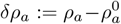 is the local deviation of *ρ*_*a*_ from its state value 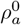, *A*^−1^ ώ 0 is the isothermal compressibility at constant strain, and *C >* 0 couples dilation with local density variation. The passive viscous stress of the elastomer is 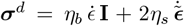, where *η*_*b,s*_ are the bulk and shear viscosities, respectively. We take the active stress ***σ***^*a*^ to be isotropic and of the form: ***σ***^*a*^ = *χ*(*ρ*_*a*_) Σ_*i*_ *ζ*_*i*_ *ρ*_*i*_ **I**, where *ζ*_*i*_ *>* 0 are activity coefficients representing contractile stress, and *χ*(*ρ*_*a*_) is a sigmoidal function which captures the dependence of the active stress on the local actin mesh density. For this permanently crosslinked actin mesh, the local density fluctuation *δρ*_*a*_ of actin is slaved to the mesh strain, 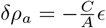 (Appendix A2).

The bound active stresslets exert contractile stresses and undergo turnover, which can in general be strain dependent; here we take the unbinding rates to be of the Bell-type form [45]: 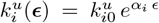, where 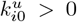, are the strain independent parts of the respective rates, and *α*_*i*_ are dimensionless numbers; *α*_*i*_ *>* 0 represent *catch bonds* where local contraction (extension) will decrease (increase) the unbinding rate of the stresslets, while *α*_*i*_ *<* 0 represent *slip bonds* where local extension (contraction) will decrease (increase) the unbinding rate of the stresslets [6–8, 35].

For a binary mixture of stresslets, the physics of segregation and the subsequent force patterning is explored by casting the hydrodynamic equations in terms of the average density *ρ* := (*ρ*_1_ + *ρ*_2_)*/*2, and relative density *ϕ* := (*ρ*_1_ − *ρ*_2_)*/*2 (*ρ*_1_ is the more contractile species):

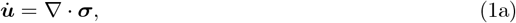

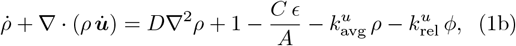

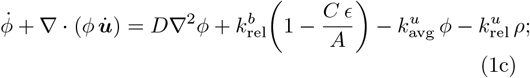

made dimensionless by setting the units of time, length and density, as 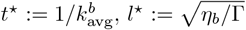 and 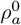, where 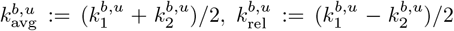 are the average and relative binding (unbinding) rates of the stresslets, respectively. The derivation of these equations together with the underlying assumptions are presented in Appendix A.

Note that despite writing the hydrodynamic equations explicitly for a binary mixture of stresslets, we may trivially use this to study the dynamics of a single myosin species by simply setting the contractility of the second species to zero (*ζ*_2_ = 0). If in addition, we set the unbinding rate 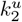 to have a slip bond nature, then species 2 can act as a *probe* molecule, highlighting regions that are depleted in myosin (species 1). [**?**]

By Taylor expanding the function *χ*(*ρ*_*a*_) about its state value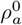, and with *δρ*_*a*_ enslaved to *ϵ*, the total stress can be recast to cubic order in strain as,

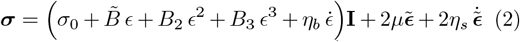

with a purely active prestress 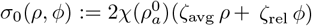, where 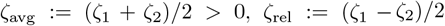 are the average and relative contractility, respectively, an activity renormalized linear elastic bulk modulus 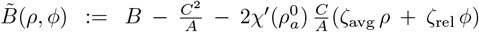 and activity generated nonlinear elastic bulk moduli *B*_2_(*ρ, ϕ*), *B*_3_(*ρ, ϕ*), linearly dependent on *ρ* and *ϕ* (see Appendix A3).

The material model, described by Eqs. (1), (2), is dynamic, renewable and active, and breaks time reversal symmetry (TRS) in that it contains terms that are not derivable from a free energy. The spatiotemporally inhomogeneous material moduli 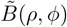 and *B*_2,3_(*ρ, ϕ*) lead to a rich effective elastic energy landscape with multiple wells, tuned dynamically by activity and material renewal. As we explore in detail later, accumulation of the stronger contractile motors (i.e., *ρ* ≫ 1, *ϕ >* 0) can make 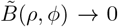 locally and even drive it to negative values, thus, triggering localized linear elastic instability, in which case the purely active higher order moduli *B*_2,3_ would stabilize the nonlinear response [4].

The renormalization of elastic moduli show up in the unusual mechanical behaviour of renewable active materials [4]. Here we focus on how these dynamic material properties drive stress patterning, directly relevant to the actomyosin cytoskeleton, leading to the emergence of singular tension structures and excitability.

## III. POSITIVE REINFORCEMENT BETWEEN ACTIVITY AND TURNOVER LEADS TO SEGREGATION OF STRESSLETS

As the first step towards force patterning, we discuss a novel segregation into domains of high and low contractility. This is readily seen from a linear stability analysis about the undeformed symmetric mixture (***u*** = **0**, *ϕ* = 0) described by the linear dynamical system, 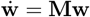 (see Appendix B2), which governs the evolution of the perturbation 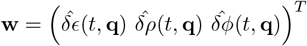 that depends on the Fourier wave vector **q**. Here, 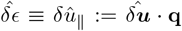 is the Fourier transform of the dilation strain field *ϵ*, and the dynamical matrix is

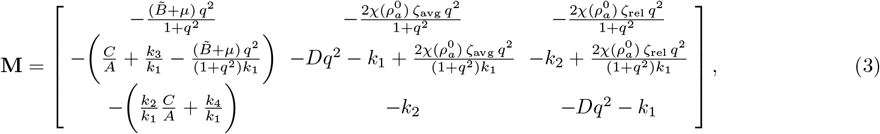

where 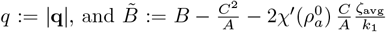 is the activity renormalized bulk modulus of the homogeneous symmetric mixture. Here the turnover rates appearing in Eq. (1) have been expanded to linear order in *ϵ*,

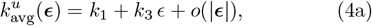

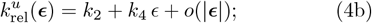

where the parameters that appear in Eq. (3), namely

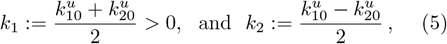

represent the average and relative bare unbinding rates (i.e., strain independent), respectively, and

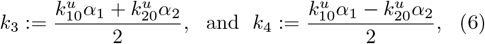

represent the coefficients of the strain dependent parts of the relative and average unbinding rates, respectively. We see that the linear dynamics of the transverse Fourier displacement mode 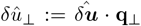 uncouples from the rest of the dynamics.

Note that **M** is non-Hermitian due to TRS breaking; as a consequence, the eigenvectors along which the perturbations propagate are no longer orthogonal to each other and may even co-align for some parameter values, as we will see later.

With the simplifying assumption that the bare rates of binding and unbinding are identical in the two myosin species, the instabilities are determined solely by the maximum eigenvalue of **M**,

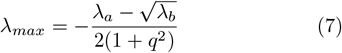

where

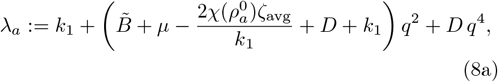

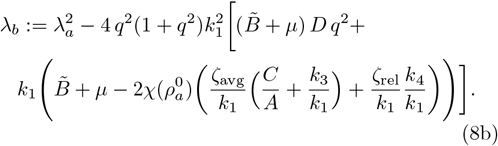

With this in hand, we are in a position to describe both the elastic and segregation instabilities of the uniform elastomer driven by active stresses and turnover (Fig. 2).

**FIG. 1.**
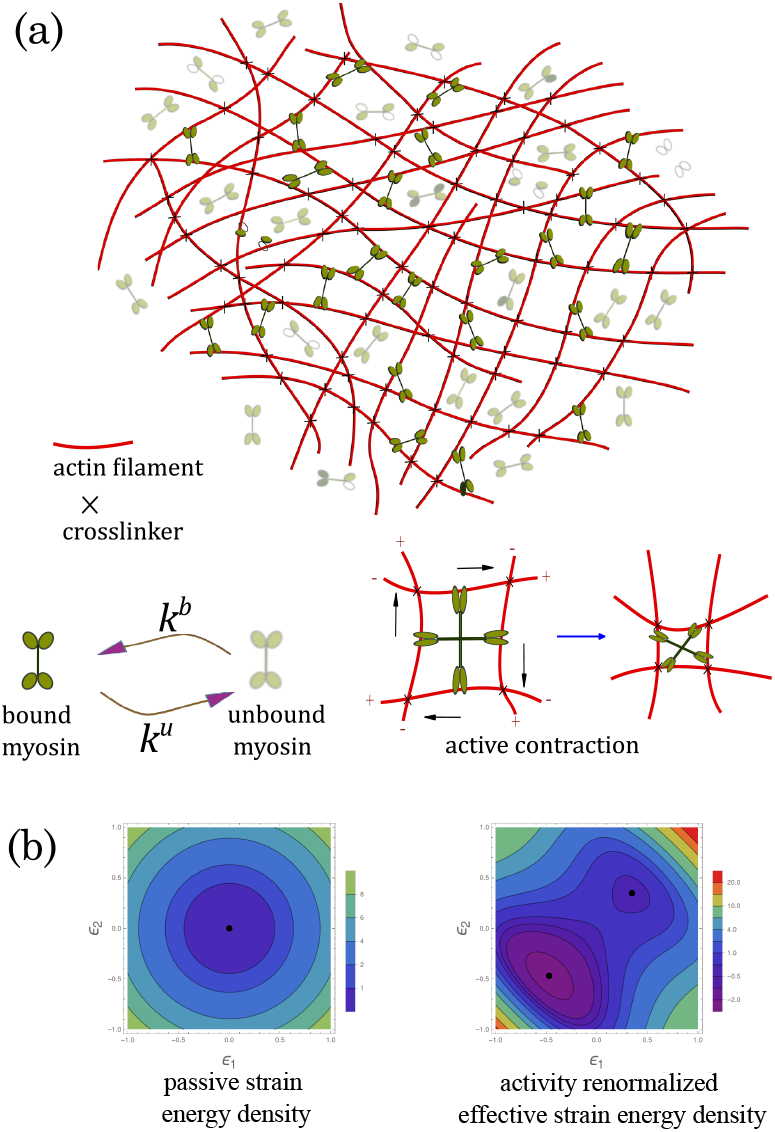
Actomyosin cytoskeleton as renewable active matter. (a) (*top*) Schematic of the actin elastomer with crosslinkers that give it rigidity, together with bound/unbound Myo-II minifilaments. (*bottom left*) Transitions from bound to unbound Myo-II and vice versa occur with rates *k*_*u*_ and *k*_*b*_, respectively. (*bottom right*) Bound Myo-II applies contractile stresses on the actin meshwork as shown. (b) (*left*) Elastic energy contour in the space of principal strains *ϵ*_1,2_ for the passive elastomer showing linear elastic behaviour with a single energy minimum (black point). (*right*) Contours of the effective elastic energy 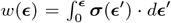 renormalized by activity. Activity generates additional minima and nonlinear stabilization [4, 35].

**FIG. 2.**
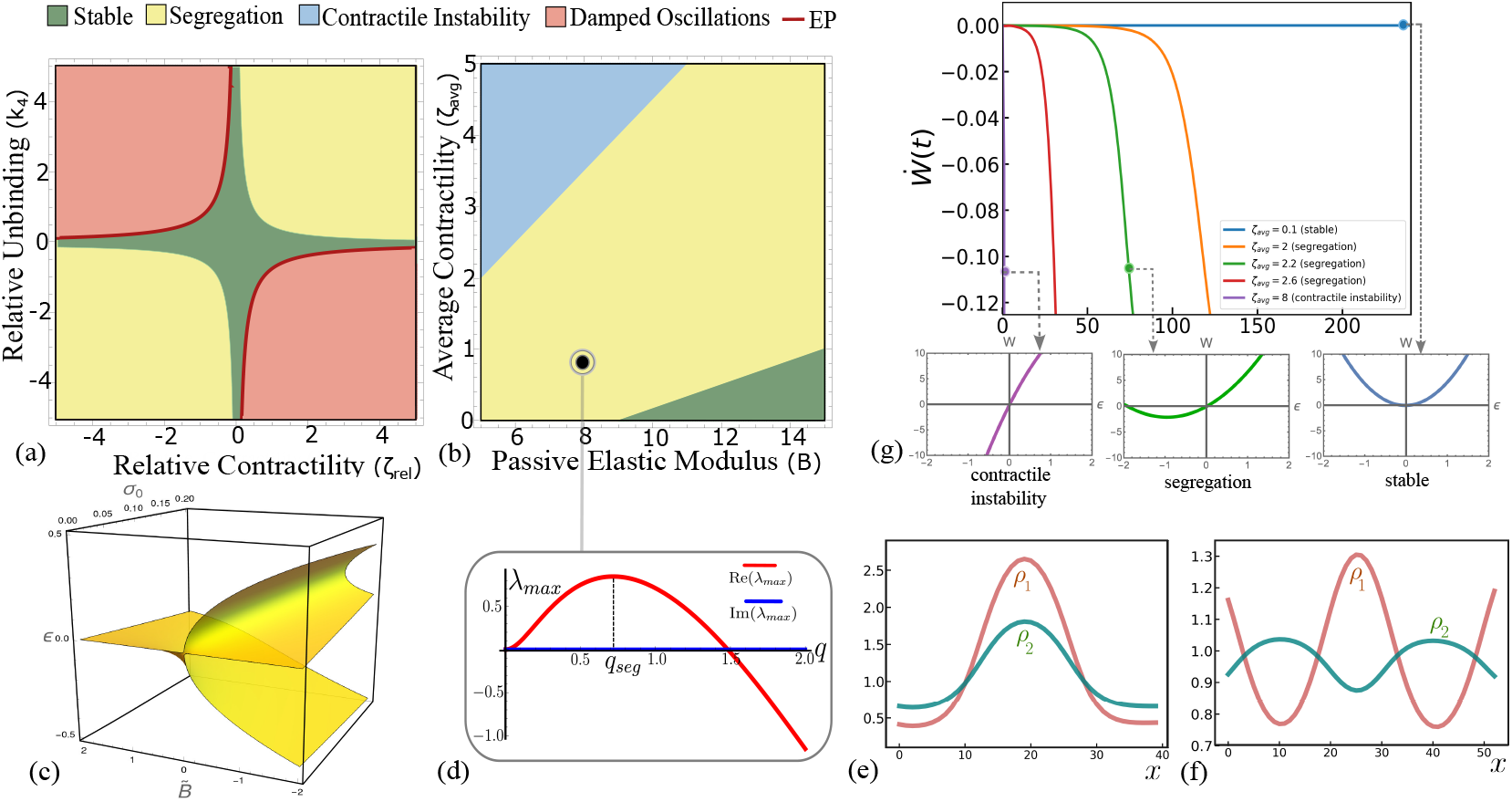
Linear stability phase diagrams and segregation instability in a binary mixture of stresslets. (a) Phase diagram in relative unbinding versus contractility, with *B* = 8, *ζ*_avg_ = 1, shows segregation instability (yellow) for *ζ*_rel_*k*_4_ *>* 0. The red line corresponds to a line of exceptional points (EP), where the eigenvectors of the dynamical matrix **M** co-align. (b) Phase diagram in bare elastic modulus versus average contractility for *ζ*_rel_*k*_4_ *>* 0, shows transitions from the stable phase to segregation followed by elastic contractile instability, with increasing *ζ*_avg_. (c) Graph of the minimum of the effective elastic energy at a fixed value of *ρ* and *ϕ*, i.e.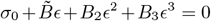, showing the elastic instability 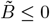 as a pitchfork bifurcation of the unstrained state into a spinodal pattern of expansion and contraction, with the active prestress *σ*_0_ acting as an “external field”. Tensile prestress *σ*_0_ *>* 0 biases the stable branch towards contraction (*ϵ <* 0). (d) Typical dispersion curve obtained from a linear analysis of the segregation instability, where *q*_*seg*_ is the fastest growing mode. (e,f) Snapshots of segregation obtained from a numerical solution of the scalar version of Eq. (1) with periodic boundary condition in 1D (see Movie 1, Movie 2). Co-localisation and separation of density peaks of the individual segregating stresselets, when (e) the two stresslets have catch bonds with *k*_3_ = 2 (see Eq. (6)), and when (f) stresslet 1(2) has catch(slip) bond with *k*_3_ = *−* 0.5. The ratio *ρ*_1_*/ρ*_2_ within the domain depends on activity, turnover and the stress jump across the domain. We have set 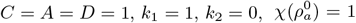 and 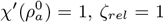 and *ζ*_*avg*_ = 2.8 throughout. (g) Power density 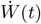 (see text) monotonically decreases in the linear segregation regime; the rate of decrease goes to −∞ as the average activity *ζ*_avg_ increases and the elastic instability is approached. Outsets show the minimum *ϵ*_*min*_ of the effective strain energy density *w*_*lin*_ at fixed (*ρ, ϕ*) moving to −∞ as *ζ*_avg_ increases (see Eq. (10)).

### Elastic (contractile) instability

Starting from a stable unstrained elastomer, we see that large enough average activity 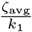 drives the renormalized longitudinal elastic modulus 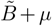 of the symmetric mixture to negative values, leading to a linear elastic instability of the underlying elastomer [4, 35] that affects all modes *q* ∈ [0, ∞) (see Fig. 2(b) and dispersion curve in Fig. B7). This elastic spinodal decomposition into patterns of positive and negative *ϵ* is a pithfork bifurcation, with prestress as an ‘external field’ controlling it (shown in Fig. 2(f)). The contracted (*ϵ <* 0) domains eventually show up as self-penetration and subsequent collapse (halted by steric effects) of the contractile mixture, unless constrained by appropriate boundary conditions [4]. The microscopic origin of this elastic instability can be attributed to the microbuckling [44], the loss of crosslinkers, resulting in relative sliding of the semiflexible actin filaments [46], to micro-wrinkling [47] and stretching-to-bending transition [48].

We will see that the force patterning that we discussed earlier is achieved through entrapping this contractile instability into segregated domains. In the linear theory, this segregation, and other nonequilibrium phases, show up in the mechanically stable regime of the active elastomer, 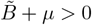.

### Segregation instability

As *λ*_*b*_ increases beyond 0, *λ*_*max*_ goes from being negative (stable) to positive, leading to a long-wavelength instability in *ϕ*. The fastest growing wave-vector *q*_*s*eg_ sets the characteristic width 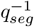 of the segregated pattern (Fig. 2(c), Eq. (C1)), provided 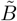 is bounded between,

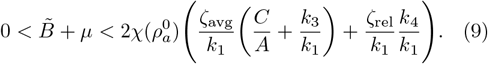

This linear segregation regime is typically realised when the relative activity *ζ*_rel_ and strain dependent unbinding *k*_4_ have the same sign, which implies 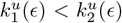, since the stresslets are contractile. To drive segregation, the stresslet with stronger contractile activity must have a lower strain-dependent unbinding rate. Note that the density peaks of the individual stresslets colocalise (Fig. 2(e), Movie 1) unlike in conventional phase separation. This is reminiscent of the co-assembly of Myosin IIA and IIB during the formation of stress fibers [15, 16, 21], and occurs when both stresslets exhibit catch-bond behaviour (*α*_1_, *α*_2_ *>* 0). For slip-bond response of the weaker species (*α*_1_ *>* 0, *α*_2_ *<* 0), the individual density peaks separate (Fig. 2(f), Movie 2).

### Driving force for linear segregation

Since the stresslets do not directly interact with each other, the driving force for this segregation in the linear theory must come from their indirect interaction through the elastomer strain. This is best seen by defining the activity renormalized “elastic energy density” in the linear elastic regime, as 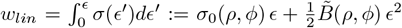. Now for any fixed value of *ρ* and *ϕ*, the minimum of *w*_*l*in_ occurs at the strain

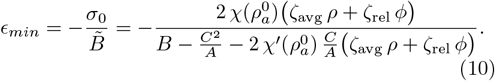

Without activity, *ϵ*_*m*in_ = 0. With activity and large *ζ*_rel_ (implying segregation), *σ*_0_ is large and 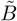 is a small positive number; hence, *ϵ*_*m*in_ becomes highly contractile. As the elastic instability is approached with further increase in activity, *ϵ*_*m*in_ → −∞ (see Fig. 2(g) insets).

Though we have not found a completely analytical proof, we see from a 1D numerical solution in a system of size *L*, that the power density 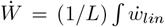 *dx* associated with linear segregation regime is negative (Fig. 2(g)), i.e., *W* is a Lyapunov functional driving segregation of the stresslets.

## IV. FINITE-TIME ELASTIC SINGULARITIES

As time passes, nonlinear terms in the dynamical equations become prominent; the growing spinodal patterns predicted by the linear analysis, start to shrink into singular structures that carry tension. This behaviour is quite unlike the usual segregating mixture, where non-linearities temper the exponential growth of the domains to a slower power-law growth [42].

To see this, let us focus on a single domain of the evolving spinodal pattern in 1D of width 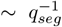 and bounded by two interfaces Γ^*±*^, Fig. 3(a). The segregation field *ϕ >* 0 inside the domain and *ϕ <* 0 outside. The forces *∂*_*x*_*σ* acting across this domain cause it to gradually shrink, concentrating the more contractile stresslet within. Eventually, the center of this shrinking domain enters the elastic instability regime 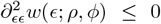 when the stresslet density is large enough (Fig. 3), following which there is no escape from collapse, leading to the formation of singular force carrying structures in finite time. These singular tensile structures which harbour the more contractile stresslet, remain well separated owing to the active prestress in the surrounding elastic medium which predominantly harbours the less contractile stresslet.

**FIG. 3.**
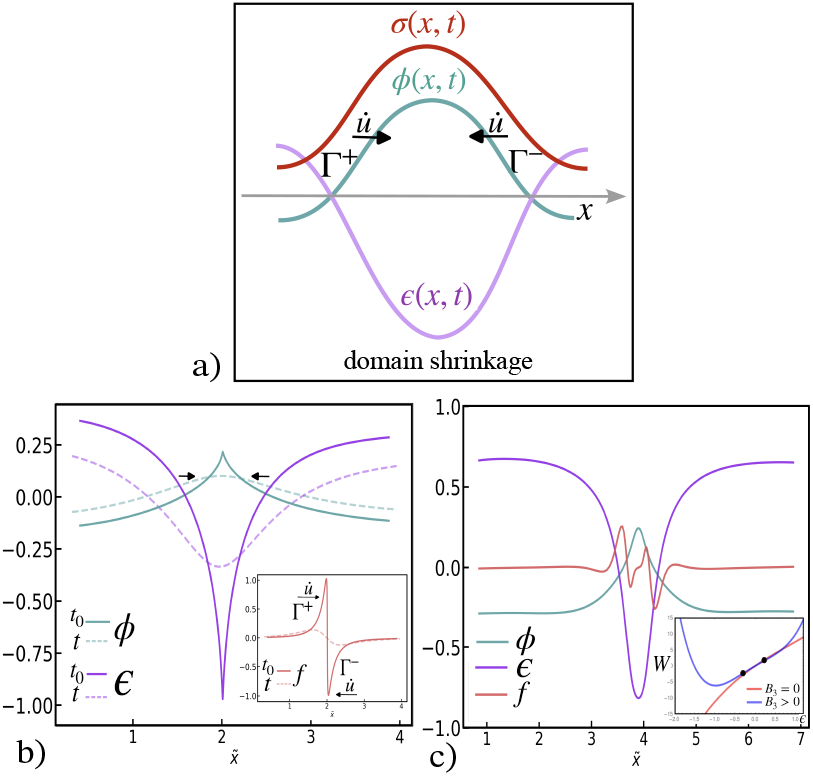
Mechanism of singularity formation. (a) Schematic of a single domain showing the profiles of the segregation field *ϕ*, strain *ϵ* and stress *σ*. The magnitude of the velocities of the interfaces Γ^*±*^ bounding the domain are 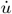, causing them to move towards each other as the singularity develops. (b) Snapshots from 1D numerical simulation of a single segregated domain showing the segregation parameter *ϕ* and strain *ϵ* profiles at an early time point *t* (dashed line) and later time point *t*_0_ (solid line) at which the singularity develops when the steric term *B*_3_ = 0. Inset shows the corresponding force 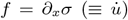 that drives the velocity of the interfaces Γ^*±*^ towards each other, leading to growing amplitudes of *ϕ* and *ϵ* and an eventual singularity. All plots are shown in the deformed coordinate 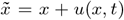. Parameters are 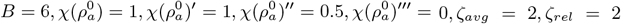. (c) Physical resolution of the singularities in *ϕ, ϵ* and *f*, by introducing a large steric term *B*_3_ = 10. (*Inset*) Comparison of the effective elastic energy landscapes for *B*_3_ = 0 and *B*_3_ = 10. The two black points 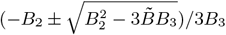, for fixed (*ρ, ϕ*), mark the elastic spinodals bounding the unstable regime of the energy landscape for a finite *B*_3_ *>* 0; the spinodals move to infinity as *B*_3_ *→* 0. See Movie 3 for (b) and (c).

In an actual physical situation, the elastic singularity is never reached and one ends up with highly concentrated tensile regions of finite width set by the constraint of steric hindrance, represented by the *ϵ*^3^ term in the effective elastic stress (Eq. (2)), see Fig. 3(c). This is consistent with the observations [15, 16] on the segregation of the more contractile non-muscle Myosin IIA from the less contractile Myosin IIB,C.

### Self-similar solution for finite-time singularity

Following the above discussion, we analyse the approach to the finite-time singularity starting from the linear segregation pattern, by dropping the steric stabilising cubic term in elastic stress and the turnover of stresslets. We also neglect the viscous stresses for convenience, though this does not pose any difficulties. The dynamical equations in 1D then reduce to the conservative system

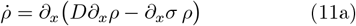

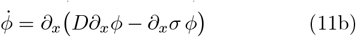

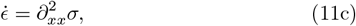

where

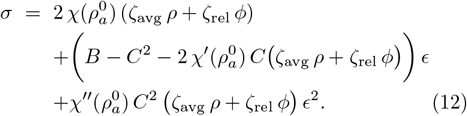

During the initial stages of linear segregation when *ρ, ϕ* and *ϵ* are relatively small, the first two terms (active prestress and linear elasticity) in the stress field Eq. (12) dominate. However, as *ρ, ϕ* and *ϵ* grow exponentially due to linear instability, they become large inside the segregated domains where local density is high. When this happens, the lowest order nonlinearity, the *ϵ*^2^ term in Eq. (12) starts dominating over other terms. This dominance of a particular power of the driving field in the final approach to the singularity, is at the origin of the scale invariance of the equation, and, hence, of the solution near the singularity [43]. Note that in the relatively low density regions between the high density domains, the dominant term is the active prestress, as the effective bulk modulus in the second term in Eq. (12) is small. Thus the interface between the high and low density regions, is *pushed* by the active prestress and *pulled* by the contractile instability.

We expect that there exists a fixed space-time location (*x*_0_, *t*_0_) at which the the first singularity appears. The scale invariance of the solution near this singularity implies that, starting from a typical segregated domain at time *t < t*_0_ as shown in Fig. 3, the height of the profiles *ρ, ϕ* and *ϵ* must grow and the domain width must shrink at *t* → *t*_0_, as power laws with appropriate *universal* scaling exponents. This a consequence of the fact that they arise solely from the structure of the partial differential equations, and are independent of initial conditions [43]. This motivates us to express the solution as a self-similar form,

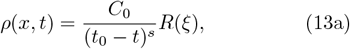

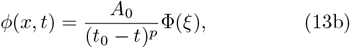

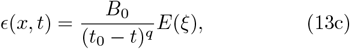

with the scaling variable, *ξ* := (*x* − *x*_0_)*/*(*t*_0_ − *t*)^*r*^. In the above, *A*_0_, *B*_0_ and *C*_0_ are constants; *s >* 0, *p >* 0, *q >* 0 and *r >* 0 are the scaling exponents; and *R*, Φ and *E* are analytic scaling functions.

With these scaling forms, we are left with the following unknowns – the blowup time and location *t*_0_, *x*_0_, the exponents *s, p, q* and *r*, and the similarity profiles *R*(*ξ*), Φ(*ξ*) and *E*(*ξ*). These are obtained by substituting the similarity forms Eqs. (13) into the Eqs. (11). Since diffusion cannot produce any finite-time singularity, we will ignore the diffusion term in the *ρ* and *ϕ*-equations.

Since we have ignored binding-unbinding, *ρ*(*x, t*) and *ϕ*(*x, t*) are conserved, thus

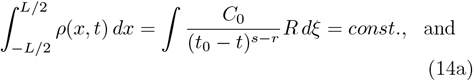

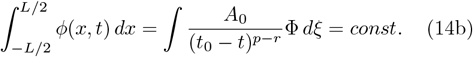

which immediately implies *s* = *p* = *r*. To get the asymptotic form of the singularity, we analyse the coupled non-linear ODEs written in terms of the scaling variable *ξ* (see Eqs. (D3), (D4) and (D5) in Appendix D) using the method of dominant balance [43]. The largest order non-linearity (coming from the *ϵ*^2^ term in Eq. (12)) should dominate near the singularity, and must balance the time derivative terms. Thus,

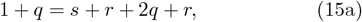

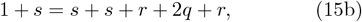

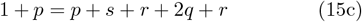

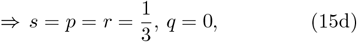

which immediately leads to the following scaling forms,

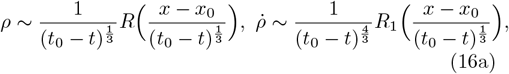

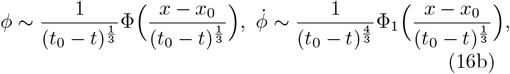

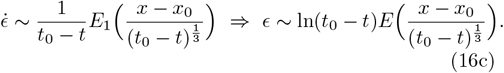

The initial data in *ϕ* is of the form 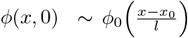, parameterized by an initial width *l*. We obtain the blowup time *t*_0_ using dimensional analysis: ∼ *τ* (*l/𝓁*)^3^, where 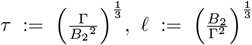, and 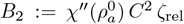. Thus the space-time location of the first blowup (*x*_0_, *t*_0_) will depend on the initial data that we determine from the numerical solution.

To obtain the scaling functions, we solve the nonlinear ODEs near the singularity

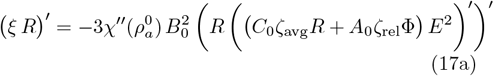

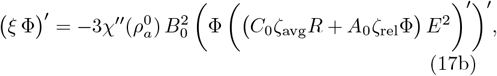

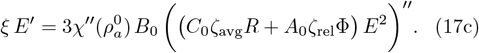

with appropriate boundary conditions. This second order system of ODEs requires two boundary conditions each. One natural choice for the boundary conditions comes from symmetry:

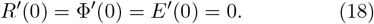

The other boundary conditions come from the asymptotics of the profiles at *ξ* → ±∞ that corresponds to the limit *t* → *t*_0_. In this limit, for *x* − *x*_0_ ≠ 0 (away from the singularity location), the singular solution must match the time independent outer solution [43]. Hence, the boundary conditions for the above ODE system is obtained by setting the time derivative terms on the left hand sides of Eq. (17) to zero:

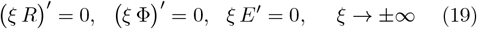

implying

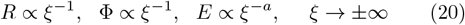

for any choice of *a* ≥ 1. The precise similarity profiles *R*, Φ and *E* can in principle be obtained by solving the non-linear ODE system Eq. (17), subject to boundary conditions Eq. (18) and Eq. (20). Note that, as the singularity is approached at *t* → *t*_0_, the strain dependent turnover term,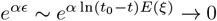. This provides an aposteriori justification for our neglect of turnover in our scaling analysis.

### Numerical solutions of the blowup

We use Dedalus pseudospectral solver [49] to solve the system of PDEs Eqs. (1) in 1D, with variables represented on a periodic Fourier basis. Periodic basis functions like the Fourier basis provide exponentially converging approximations to smooth functions. The fast Fourier transform (implemented using sciPy and FFTW libraries [49]) enables computations requiring both the series coefficients and grid values to be performed efficiently. For time evolving the system of equations, we use a SBDF2 time-stepper, a 2nd-order semi-implicit Backward Differentiation formula scheme [50]. Our initial conditions are a homogeneous unstrained symmetric mixture *ϕ* = 0, to which we add noise sampled from a uniform distribution between [− 0.1, 0.1]. We use 500 modes (*N* = 500) in a domain length of *L* = 35 and a time-step (*dt*) of 10^*−*4^ to solve Eqs. (1). We use the following set of parameters: 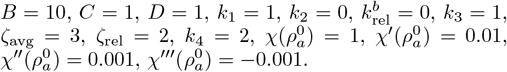.

On solving we find that after the initial linear stability regime, singularities in *ρ, ϕ* and *ϵ* develop in regions of high myosin density. The profiles of *ϕ* and *ϵ* near the singularity, at different time points, are shown in Fig. 4(a) and Fig. 4(c), respectively. We numerically determine the first blowup time *t*_0_ and location *x*_0_, from the divergence of the maximum (minimum) value of *ϕ* (*ϵ*), as seen in the insets of Fig. 4(a), (c). In agreement with our scaling analysis, we see an excellent collapse of the data for the scaled profiles of *ϕ* and *ϵ*, see Figs. 4(b), (d). This agreement between the numerical solutions of Eqs. (1) and the scaling analysis of Eqs. (11) reaffirms the irrelevance of myosin turnover in the neighbourhood of the singularity.

**FIG. 4.**
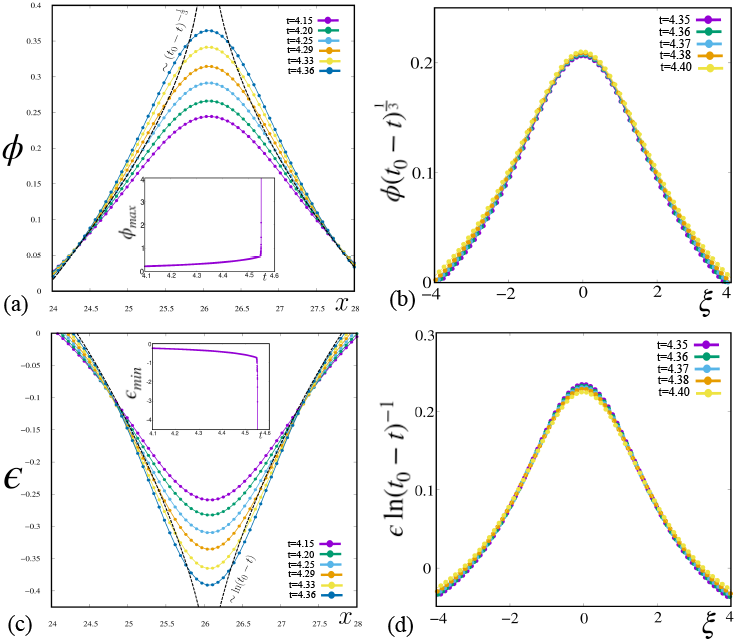
Numerical analysis of finite time singularities. (a)-(d) Numerical results verifying the scaling profiles of *ϕ* and *ϵ* and the formation of a singularity in finite time. (b,d) show scaling collapse of the *ϕ* and *ϵ* profiles near the singularity, as predicted from theory; 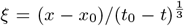 is the scaling variable. Starting from a homogeneous unstrained symmetric state, a 1D numerical solution of Eq. (1) gives the value of the finite time blowup *t*_0_ = 4.55 and blowup location *x*_0_ = 26.06 (see insets in (a) and (c)).

Since our numerical method, does not allow us to approach the singular point arbitrarily closely, we use our numerics to test the asymptotic convergence of the profiles approaching the singularity. We employ the Cauchy-convergence criterion for the sequence *ϕ*(*x, t*_*n*_), for a fixed value of *x* near *x*_0_ and where the sequence {*t*_*n*_} steadily approaches *t*_0_. In Fig. E9, we have plotted *ϕ*(*x, t*_*n*+1_) − *ϕ*(*x, t*_*n*_) as a function of *t*_*n*_, at three spatial locations *x* − *x*_0_ = 0, *x* − *x*_0_ = 0.6, and *x* − *x*_0_ = 1. We see that *ϕ*(*x, t*_*n*+1_) − *ϕ*(*x, t*_*n*_) → 0 faster for points closer to the centre of the profile; the convergence rate is maximum at *ϕ*_*max*_.

These singularities are *physical* in that their resolution involves incorporation of additional physical effects such as steric hindrance (represented by the *ϵ*^4^ term in *w*). With this, the singularity is never reached, resulting in highly concentrated tensile regions of finite width, which could be taken to be the size of non-muscle myosin-II bipolar filaments [13].

The appearance of finite time elastic singularities that lead to zones of high concentration of both the actin meshwork and myosin stresslets is reminiscent of caustics discussed in the context of the Zel’dovich model for the formation of large scale self-gravitating structures of the universe [51] and verified in extensive numerical simulations, such as in [52]. We observe that at the blowup location *x* = *x*_0_ where *ϵ* ≈ − 1 (Fig. 3(c)), actin meshwork density in the deformed or Eulerian configuration 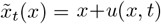, given by 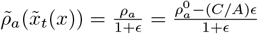, goes to + ∞ as *t* → *t*_0_. Moreover, this meshwork density blowup occurs before the stresslet density blowup, as seen both from the similarity forms in Eq. (16) which shows that the strain rate 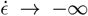 faster than *ρ*_*i*_ → ∞, at the same spatial blowup location *x*_0_, and from the numerical solution (see Fig. E8). These *elastic caustics* as singularities of Lagrange maps 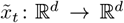 in higher dimensions will appear as more general Lagrangian catastrophes [51, 53], of a variety of co-dimensions (points, lines, surfaces etc.) and topology, where the associated deformation map 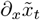 degenerates.

We note from our 1D numerical analysis, that the maximum strain magnitude within singular structures is of order 0.8 while outside it can reach a value of 2.5, suggesting that geometrical nonlinearities are significant. This will be particularly relevant in 2D.

## V. MOVING, MERGING AND COARSENING OF SINGULARITIES

In the above scaling analysis, we have assumed that the singular structures are described by symmetric profiles of the fields, which lead to static finite-time singularities. However, our numerical analysis shows that the singular structures, tempered by steric hindrance, can move. Such *moving* singularities must be associated with an asymmetry in its density profile and discontinuities in bulk stress and mass flux across it. Indeed moving singularities show up as moving stress fibers whose interactions and coalescence have been reported in *in vivo* studies [5].

Before proceeding, we draw attention to the study of the *nucleation* instability of clusters of bound myosin [41], as opposed to the *spinodal* instability studied here. We show that on nucleating a symmetric myosin pulse of finite width, the pulse grows if the amplitude is beyond a threshold, else it relaxes to a uniform background. Further, on nucleating an asymmetric pulse, a large enough asymmetry drives a translational instability. Using a boundary layer analysis, we derive analytical forms for the asymmetric myosin density and strain profile and its velocity [41]. This information will be useful in the analysis presented below.

### Isolated moving singularity

The moving singularity at *x*_0_(*t*) carries with it the singular component of all the bulk fields, accompanied by jumps across *x*_0_(*t*). We denote the jump of the limiting values (i.e., difference between right and left) of bulk fields at *x*_0_(*t*) ⟦⟧ by and their average by ⟨⟩. Continuity of the bulk displacement field ⟦*u*⟧ (i.e., *u* = 0) implies the velocity compatibility condition [54], a kinematic requirement:

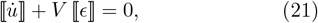

where 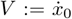 denotes material velocity of the singular point (see Appendix E). The spatial velocity of the singularity is then defined by 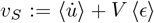. The force and mass balance equations at the singularity can be obtained, using the divergence and transport theorems for discontinuous fields, as

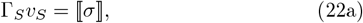

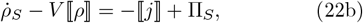

where Γ_*S*_ is the friction coefficient at the singularity, *ρ*_*S*_ is the stresslet density at the singularity, 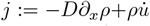 is the bulk mass flux and Π_*S*_ is the (strain dependent) singular turnover term (see Appendix E). Hence, the jumps in stress, mass density and mass flux across each singularity act as the driving force for movement of the singularity and its height variation.

### Interacting moving singularities

When the number density of singularities is high, they will interact with each other via the long-range elastic strain. Since the moving singularities are associated with an asymmetric density profile, there could be additional non-reciprocal interactions between them, possibly leading to dynamical phenomena discussed in [55, 56].

### Merging and Coarsening

Thus far we have established that following the initial segregation and finite time collapse, we are generically left with a distribution of moving and interacting singularities. Subsequent evolution involves the merger and subsequent coarsening of singularities. There are two important features of this coarsening dynamics that emerge from our 1D analysis – (i) because of the long range strain, and velocity compatibility Eq. (21), the analysis of the dynamics of merger needs to be global, and (ii) coarsening proceeds so as to make the stress homogeneous, i.e. *σ* = *const*., except at isolated points associated with the singularities. This is reflected in the time dependence of ∫ (*∂*_*x*_*σ*)^2^ *dx* (see Fig. 6), which shows an overall decrease in time, except at isolated time points associated with the onset of singularity mergers, a kind of Lyapunov functional.

**FIG. 5.**
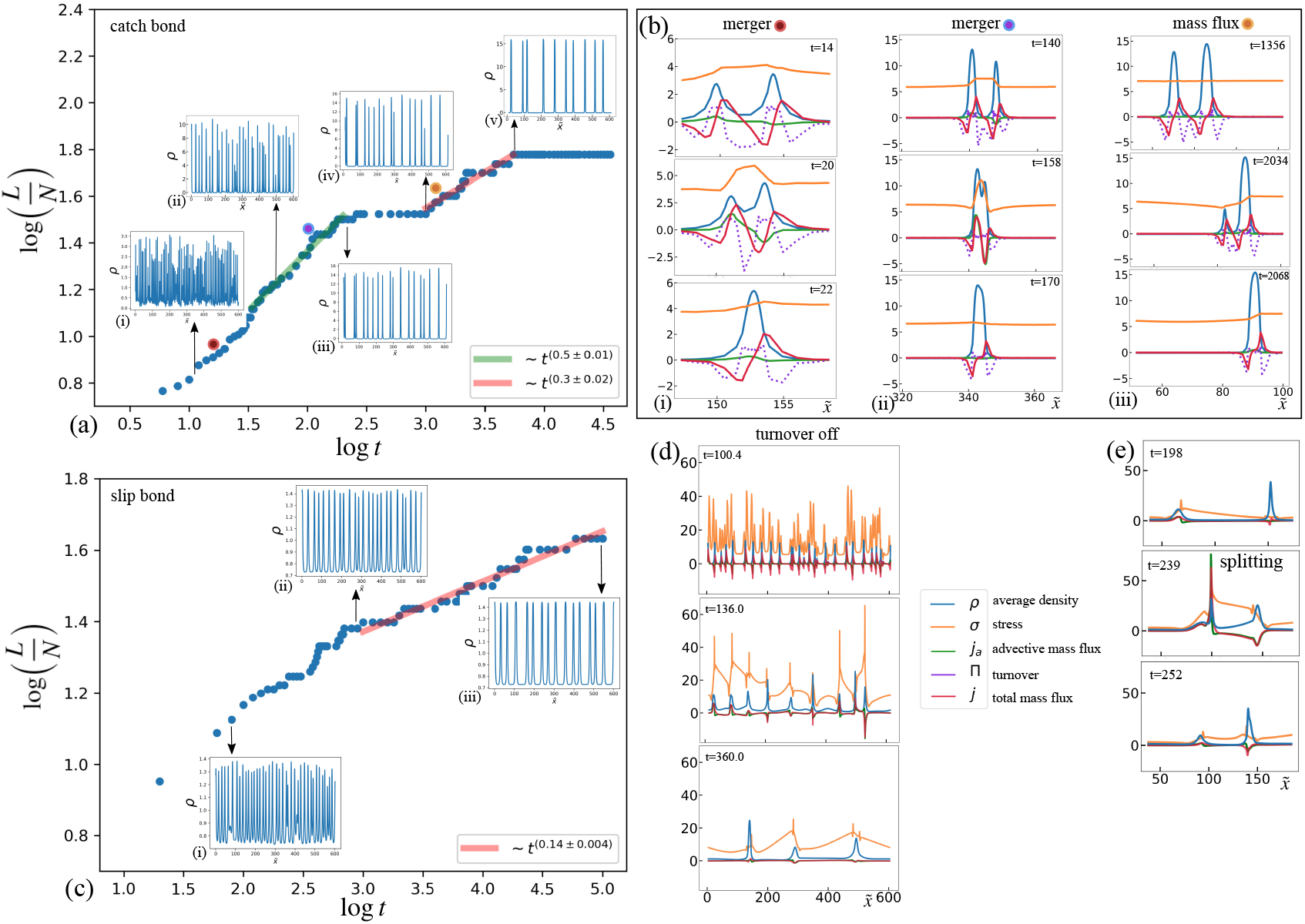
Merging of singular structures and coarsening dynamics. In figure (a) we see coarsening dynamics for a system with catch bond. The system of equations are numerically solved for the following set of parameters :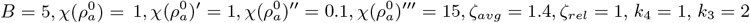. Panel (b) describes the dynamics of merger events driven by stress gradients and mass flux via snapshots from the numerical simulation described in (a). Panel (c) captures the coarsening dynamics for a system with slip bond. All the parameters remain unchanged apart from *k*_3_ = *−* 0.5. Panel (d) shows how the configurations generated in the numerical simulation in (a) at *t* = 100, evolve after the turnover is switched off. The jump in total flux ⟦*j*⟧ and stress ⟦*σ*⟧ is much higher in the absence of turnover and the merger events proceed faster as seen from the snapshots. (e) Sequence of snapshots showing splitting of singular structures after the turnover is switched off (see Movie 5).

**FIG. 6.**
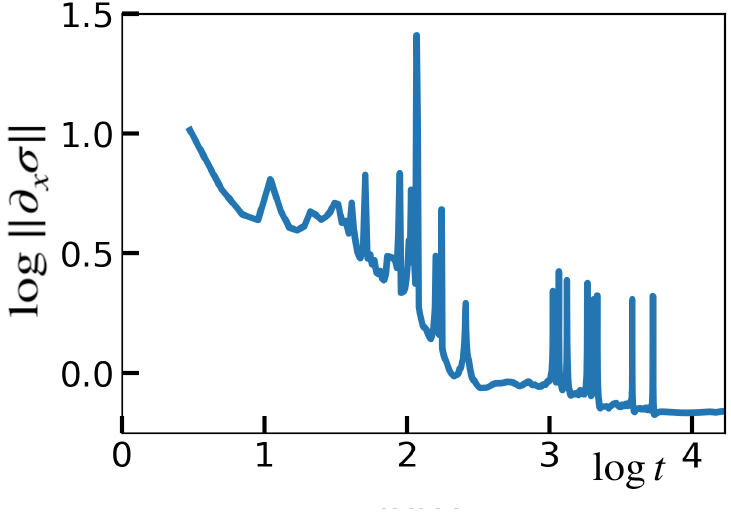
Log of *L*^2^ norm of 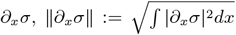, is plotted as a function of log time, in the case where the two myosin species have catch bonds (with parameters same as Fig. 5(a), Movie S4). The stress gradient shows an overall decrease in time, except at isolated time points associated with the onset of singularity mergers. This could be interpreted as a Lyapunov functional, if smoothened with a suitable mollifier.

We see that the inverse density of singularities *L/N* which measures the typical distance between singularities, grows initially and then saturates, see Fig. 5(a) and (c), for the cases where both the species are of catch and slip type, respectively. The power law growth associated with early time coarsening is indicated in the figures. The saturation at late times, either indicate arrested coarsening or at most a slow logarithmic growth indicative of activated dynamics.

To understand the dynamics of coarsening, we resort to the singular balance equations (22) for isolated singularities, and monitor the time dependence of the local density fields, stresses and currents. Soon after the establishment of the distribution of singularities, the number density of singular structures is high and the variation in the heights of the singular density profiles 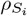 is large; here, the subscript *i* labels the singular peaks (Fig. 5(a), inset (i), and Movie 4). As a result all the driving jump terms ⟦*ρ*⟧_*i*_, ⟦*j*⟧_*i*_ and ⟦*σ*⟧_*i*_, across each singularity *i*, are large, leading to a large speed *V*_*i*_ and large mass flux 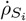 out of/into the singularities (Fig. 5(b)(i)). This drives the early mergers, since the elastic energy barrier between neighbouring singularities are easily overcome, and results in fast coarsening with a rate *t*^0.5^ (Fig. 5(a), green line). After several merger events, the singularities get sparser, the variation in the heights of the singularities gets smaller (Fig. 5(a), insets (ii,iii,iv)), the jumps in the density fields and the mass flux across the singularities become smaller – and as a result, the elastic stress gets more homogeneous (Fig. 5(b)(ii)). Together, this leads to a slower coarsening with a rate *t*^0.3^ (Fig. 5(a), pink line), associated with a driving force that mainly comes from the advective current flowing from slightly smaller singularities to higher ones, (and to a lesser extent through the turnover of myosin), see Fig. 5(b)(iii). We note that the fitted growth exponents reported here represent intermediate time dynamics, and are not universal. For *k*_3_ *<* 0, i.e., slip bond response, the stress becomes homogeneous relatively quickly, and the coarsening gets much slower with a rate *t*^0.14^ (Fig. 5(c) and Movie 6).

Eventually the variation in the heights of the singularity profiles vanishes, and the dynamics reaches a steady state, thus, arresting the coarsening, with a distinct distribution of singular structures, see the plateau in Fig. 5(a) and inset (v).

Interestingly, we find that myosin turnover, is crucial in sustaining this dynamics of singular structures over a long time. Thus, if we switch off the turnover during this coarsening stage, say at *t* = 100 in Fig. 5(a), we find a large increase in the mass flux from diffusion and advection, causing the singular structures to spread and overlap, leading to merger and faster coarsening, see Fig. 5(d) and Movie 5. We also observe the splitting of singularities, see Fig. 5(e) and Movie 5, which is reminiscent of stress fiber splitting reported in [5, 57]. This quickly leads to a single domain with high myosin density, a contractile collapse, as predicted by the phase diagram when the turnover rates are set to zero. In vivo, this eventuality can be interrupted by anchoring on other extended structures such as on microtubules or on the plasma membrane or internal membranes.

This suggests that material renewability acts as a control for the coarsening dynamics, maintaining the distribution and organisation of the singular force structures over long time scales.

## VI. DISCUSSION

Renewable active matter is a new class of active materials that exhibit novel mechanical properties such as elastic marginality, force chains, mechanical fragility and spontaneous excitability [4, 35, 41]. In this work, we have studied how the interplay between activity and material renewability can give rise to unusual segregation and nonequilibrium force patterning in living materials. A striking example is the active cytoskeleton, where a variety of myosin species—as discussed in the Introduction— act as molecular force generators and sensors, giving rise to singular extended structures carrying tension (the stress fibers).

We derive hydrodynamic equations in terms of the local actin mesh displacement and the densities of two types of myosin species that differ in their contractility and turnover rates, but which otherwise do not interact with each other. Despite this, the two myosin species undergo a spinodal segregation instability, with myosin density patterns that depend on the nature of the strain-dependent unbinding, namely, whether they have catch or slip bond response.

Subsequently the spinodal patterns become unstable due to a nonlinear coupling between the linear segregation and contractile instability. This drives domain shrinkage, leading to a finite-time singularity where the density and elastic fields display scaling behaviour. We formally associate the distribution of these singular structures with caustics. These singular structures or caustics move, merge and coarsen over time. The rate of coarsening gets slower as time progresses and eventually gets arrested. This segregation of myosin species into singular structures immediately implies spatiotemporal force patterning that spans across the system. Importantly, *material renewability in the form of myosin turnover, is crucial in maintaining this dynamics and eventual arrest of singular structures, and thus force patterning*.

We will, in another publication, extend our analysis to a fully tensorial elasticity in two dimensions; while this extension is nontrivial, the basic message of segregation followed by a finite time collapse survives. However, we note that we would need to include active boundary anchoring at the cell surface in the form of focal adhesions and adherent junctions, whose strength and positioning will select specific bulk force patterns [58, 59]. This will allow us to make direct quantitative comparisons to the dynamical patterning of stress fibers, such as those observed in experiments in cells [60], in cell extracts on micropatterned substrates [5, 61], and in vitro reconstituted systems with controlled mechanical constraints [62].

From an evolutionary materials perspective, our work suggests that neutral variations in the space of myosin binding/unbinding affinities and contractility, may facilitate favourable phenotypes [63], such as, differences in cellular localisation and the patterning of a network of forces across the scale of the cell. Further, in the context of *physical learning* [64], our work provides insights on active materials synthesis whose components learn parameter values that drive it towards specific material response, such as mechanical adaptation [38] and tensegrity [65].

We conclude by reflecting on the broader implications of material renewability in active systems—particularly in relation to material evolution and physical learning [64]. While active elastomers composed of a single myosin species can display pattern formation and contractile segregation, introducing multiple myosin species with distinct contractility and strain dependent turnover rates not only greatly enhances the richness of the emergent dynamics, but also provides a new conceptual framework for understanding how self-organizing materials can evolve toward mechanical adaptation. The wide range of contractility and binding-unbinding kinetics found in living systems equips the material to explore an extended landscape of mechanical responses (i.e., material moduli). This enables the system to self-tune [66] and evolve toward mechanically adaptive states with different functional phenotypes [4].

## VII. ACKNOWLEDGEMENTS

We thank members of the Simons Centre at NCBS for clarifying discussions. We acknowledge support from the Department of Atomic Energy (India) under project no. RTI4006, the Simons Foundation (Grant No. 287975) and the computational facilities at NCBS. MR acknowledges Department of Science and Technology (India), for a JC Bose Fellowship (SERB File No.JCB/2018/00030).

## Appendix A: Hydrodynamic Equations for a Mixture of Contractile Stresslets on an Elastomer

We describe the dynamics of active stress propagation in the active medium of the cell using hydrodynamic equations for the crosslinked actin mesh and the density of different species of myosin filaments, embedded in the viscous cytosol.

### A1. Equations for the Crosslinked Actin Meshwork embedded in the Cytosol

We start with a passive 2-dimensional elastomeric meshwork of mass density *ρ*_*a*_, whose displacement with respect to an unstrained reference state is ***u***. The meshwork moves in the fluidic cytosol whose velocity is ***v***. The hydrodynamic equations for its linear momentum balance and mass balance are, therefore,

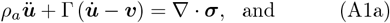

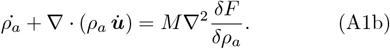

Here, Γ *>* 0 is the friction coefficient of the elastomer with respect to the fluidic cytosol, and ***σ*** is the total stress in the elastomer; *M* represents the mobility of permeation of the meshwork. *F* is the free energy functional for the passive meshwork:

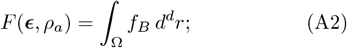

where the free energy density *f*_*B*_(***ϵ***, *ρ*_*a*_) depends on the linearized strain ***ϵ*** := (∇ ***u*** + ∇ ***u***^*T*^)*/*2 of the elastomer, and the mass density *ρ*_*a*_.

The hydrodynamic equations of the fluidic cytosol are given by

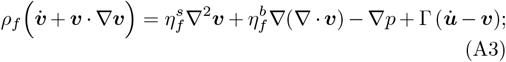

here, *ρ*_*f*_ is the density of the fluid, and 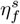 and 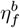 are the shear and bulk viscosities of the fluid. The fluid pressure *p* appears due to the total incompressibility of the meshwork-fluid system:

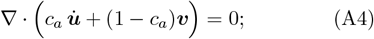

here, *c*_*a*_ is volume fraction of the meshwork. Here, for convenience, we ignore the hydrodynamics of the fluid, permissible in the limit when *c*_*a*_ ≈ 1.

### A2. Equations for the Stresslets

Consider a mixture of active contractile stresslets with different contractilities undergoing turnover onto this elastomer. We assume that the stresslets binding onto the elastomer are recruited from an infinite pool of stresslets unbound to the elastomer, and, hence, disregard the dynamics of these unbound stresslets. Restricting to a binary mixture, let *ρ*_*i*_, *i* = 1, 2, be the density fields of the two species of bound stresslets. The bound stresslets get advected by the local velocity 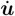 of the elastomer, and diffuse on it with the same diffusion coefficient *D*. Let the stresslets bind onto the elastomer with rates 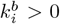, and unbind from the elastomer with rates 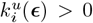 that in principle depends on the strain ***ϵ*** of the elastomer. The dynamics of these bound stresslets is, hence, governed by

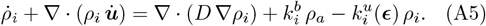

We will assume the Hill form for the unbinding rates: 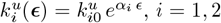 where 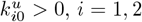, are the strain independent parts of the respective rates, *ϵ* := tr ***ϵ*** is the dilational strain, and the dimensionless numbers *α*_*i*_ capture whether the bond is catch or slip type: *α*_*i*_ *>* 0 ensures that local contraction (extension) will decrease (increase) the unbinding of the stresslets, i.e., the bond is of catch type, while *α*_*i*_ *<* 0 ensures that local contraction (extension) will increase (decrease) the unbinding of the stresslets, i.e., the bond is of slip type.

We consider the actin elastomer to be permanently crosslinked and that the elastomer density is slaved to the strain, i.e.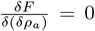. Thus, *δρ*_*a*_ ∝ −*ϵ* (obtained from a variation of (A6) below), where 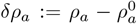 is the deviation of the elastomer density from its state value 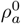.

### A3. Constitutive Equations

The elastic stress 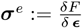 comes from the free-energy functional *F* (***ϵ***, *ρ*_*a*_) = ∫*d*^2^*rf*_*B*_ of an isotropic linear elastic medium, where

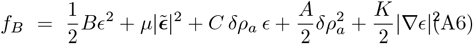

is the free energy density; with the elastic moduli *B, µ >* 0, and *C >* 0, *A >* 0 from thermodynamic stability. The isotropic elastic stress is

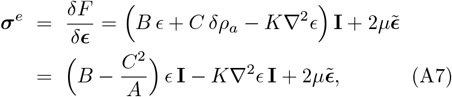

noting that 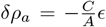. The elastic modulus *B* of the actin mesh is set by the crosslinker density. The passive viscous stress of the elastomer is 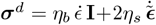, where *η*_*b,s*_ are the bulk and shear viscosities, respectively.

At the macroscopic/coarse-grained scale, the isotropic active stress ***σ***^*a*^ is of the form ***σ***^*a*^ = Δ*µ χ*({*ρ*_*a*_})*ζ*(*ρ*_*i*_)**I**, where Δ*µ* is the chemical potential change due to ATP hydrolysis, and *χ*(*ρ*_*a*_) is a sigmoidal function that encodes the dependence of the active stress on the meshwork density. We will take Δ*µ* = 1 for simplicity.

We assume that the stresslets do not interact directly but only via the strain of the elastomer. Hence, the function *ζ*({*ρ*_*i*_}) can be additively decomposed into individual contributions coming from each stresslet species, i.e., *ζ*({*ρ*_*i*_}) = Σ_*i*_ *ζ*_*i*_(*ρ*_*i*_). We assume the functions *ζ*_*i*_(*ρ*_*i*_) to be linear in *ρ*_*i*_, i.e., *ζ*_*i*_(*ρ*_*i*_) = *ζ*_*i*_*ρ*_*i*_ (no sum over *i*) where the contractilities *ζ*_*i*_ *>* 0 are different for different species *i*. For a binary mixture, Taylor expanding the function *χ*(*ρ*_*a*_) about the state value 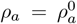 upto cubic order leads to

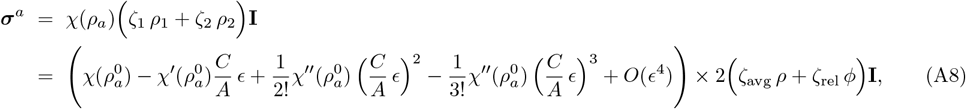

where we have introduced the the average and the relative densities of the bound stresslets:

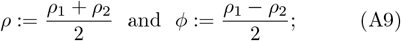

and

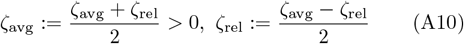

as the average and relative contractility, respectively. Note that *ϕ* is the order parameter for segregation.

### A4. Non-dimensional Governing Equations

We will further assume the overdamped limit of the elastomer, i.e., 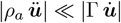, and introduce

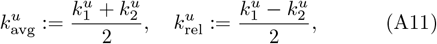

the average and relative unbinding rates, respectively;

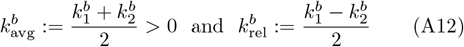

the average and relative binding rates, respectively. Taylor expanding the Hill form of unbinding, we write

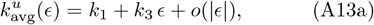

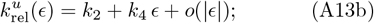

where

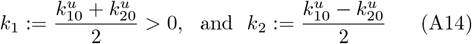

are the bare (i.e., strain independent) average and relative unbinding rates, respectively, and

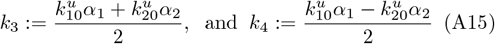

are the coefficients of the linear strain dependent parts of the relative and average unbinding rates, respectively.

Note that if *ζ*_rel_ = 0, *k*_2_ = 0 and *k*_4_ = 0, the distinction between the two contractile species disappears and the system becomes effectively one species. Hence, these three parameters in our model cannot be made zero simultaneously.

Let the characteristic time scale be 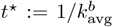, the characteristic length scale 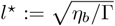, and the characteristic density 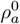. We non-dimensionalize all the variables with the following redefinitions:

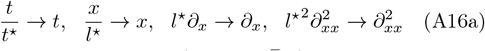

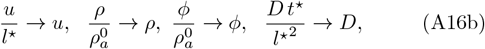

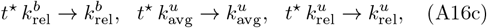

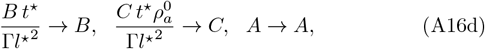

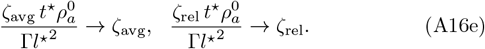

With this, the non-dimensional form of the governing equations become

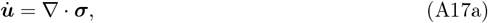

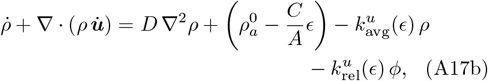

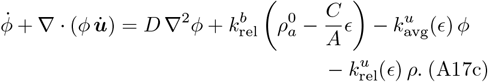

with

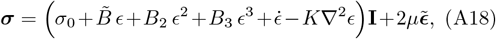

as the non-dimensional stress, where

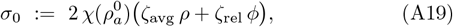

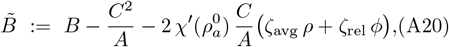

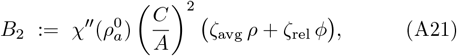

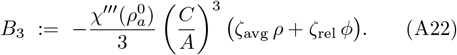

Here, *σ*_0_ is the purely active back pressure, 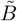 is the activity renormalized bulk modulus of linear elasticity, *B*_2_ and *B*_3_ are purely active nonlinear bulk moduli, which depend on *ρ*_1_ and *ρ*_2_. For the effective material to show contractile response, 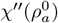 and hence *B*_2_ needs to be positive. For stability of the nonlinear elastic material, 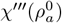 must be negative (rendering *B*_3_ positive).

## Appendix B: Linear Stability Analysis

### B1. Homogeneous unstrained steady state

For the homogeneous unstrained steady states of the system (A17), we have ***u*** = **0**, and ∇*ρ* = ∇*ϕ* = **0**. The dynamical equations reduce to

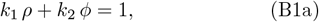

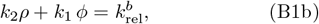

which yields

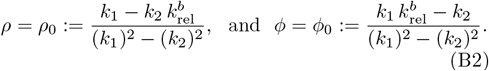

For *ρ*_0_ *>* 0, we require either 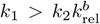 and 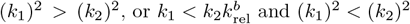.

### B2. Linear stability of the homogeneous unstrained steady state

To study the linearized dynamics of (A17) around the homogeneous unstrained steady state, we substitute ***u*** = *δ****u***, *ρ* = *ρ*_0_ + *δρ* and *ϕ* = *ϕ*_0_ + *δϕ* in (A17), neglect all the terms containing higher powers of *δ****u***, *δρ* and *δϕ*, and obtain

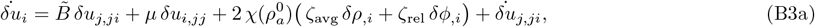

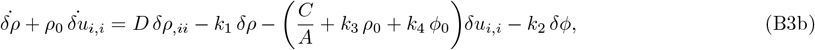

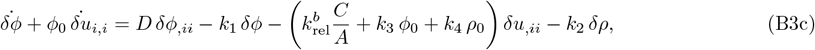

where 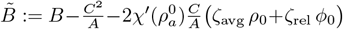 is now the (spatiotermporally constant) renormalized linear elastic modulus at the homogeneous steady state.

The Fourier transform of (B3) with respect to **x** is obtained by substituting the anstatz 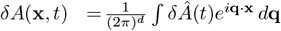, where *A* stands for ***u***, *ρ, ϕ*, and **q** is the wave vector, with *q* := |**q**|. We rewrite the resulting equations in terms of dilation mode 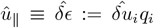, the Fourier transform of *ϵ* = div ***u***. We do not write the dynamics of the shear mode 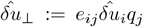 here as it uncouples from the rest of the dynamics (*e*_*i*j_ denotes the 2D Levi-Civita symbol, with *e*_11_ = *e*_22_ = 0 and *e*_12_ = −*e*_21_ = 1). The final linear equations (assuming the interface term *K* = 0) are

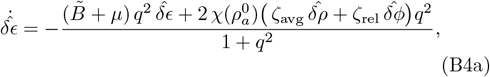

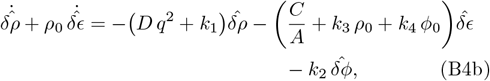

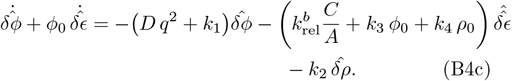

We focus on the special case *ϕ*_0_ = 0, i.e., symmetric binary mixture. From (B2), we obtain the necessary condition to maintain this, namely, 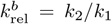. As a consequence we see that *ρ*_0_ = 1*/k*_1_, and 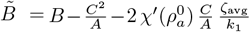. The resulting dynamical system in the Fourier space can be written as 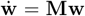, where 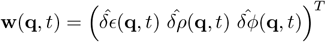, and

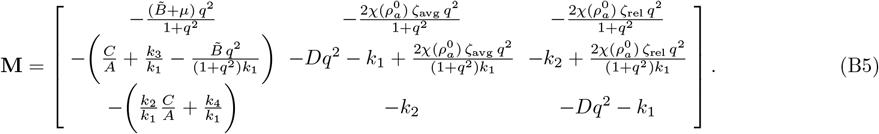

If *λ*_*i*_(*q*) are distinct eigenvalues of **M**(*q*) and **v**_*i*_(*q*) are the corresponding (linearly independent) eigenvectors, then we can write the general solution as

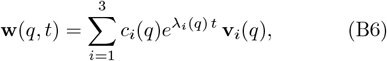

where the coefficients *c*_*i*_(*q*) are the projections of the initial data **w**(*q*, 0) along the respective eigenvectors: (*c*_1_(*q*) *c*_2_(*q*) *c*_3_(*q*))^*T*^= **V**(*q*)^*−*1^**w**(*q*, 0), where **V**(*q*) := **[v**_1_(*q*) **v**_2_(*q*) **v**_3_(*q*)] is the matrix containing the eigen-vectors as columns.

The three eigenvalues, assuming *A* = 1, are *λ*_1_ = −*k*_1_− *D q*^2^ and *λ* 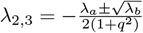, where

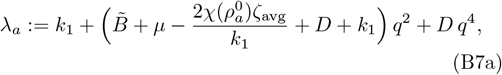

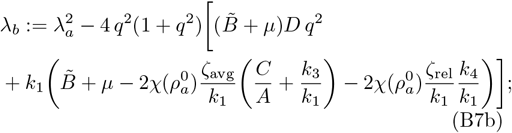

and the corresponding eigenvectors are

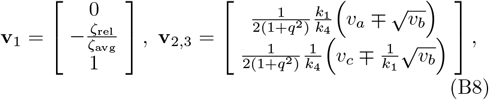

where *v*_*a*,b,c,d_ have rather long expressions that we do not write here. Since the matrix **M** is non-Hermitian (due to presence of activity and turnover), its eigenvalues *λ*_*i*_ are not real and the eigenvectors **v**_*i*_ are not orthogonal in general. We make one further simplifying assumption: 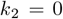, hence, 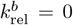, meaning that the bare (strain independent) part of the unbinding rates are idential (i.e., 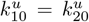, and the binding rates are identical as well (i.e.,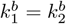). It follows that 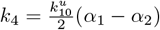.

Note that, (a) the coupling *M*_12_ between 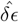 and 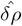 in the 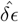 -equation and the coupling *M*_21_ between 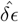 and 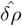 in the 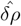-equation can be of opposite signs depending upon the relative signs and magnitudes of *k*_3_ and 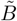; (b) the coupling *M*_13_ between 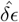 and 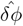 in the 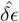 - equation and the coupling *M*_31_ between 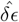 and 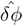 in the 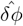 -equation are of opposite signs when *ζ*_rel_ and *k*_4_ have the opposite signs; and (c) the coupling *M*_23_ between 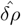 and 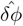in the 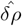 -equation can have any sign while the coupling *M*_32_ between 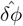 and 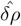 in the 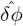 -equation is zero. Hence, 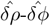 interaction is always non-reciprocal, while the 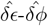 interaction is non-reciprocal when the stronger stresslet unbinds faster; 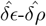 interaction can be non-reciprocal depending upon the relative signs and magnitudes of *k*_3_ and 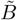.

**FIG. B7.**
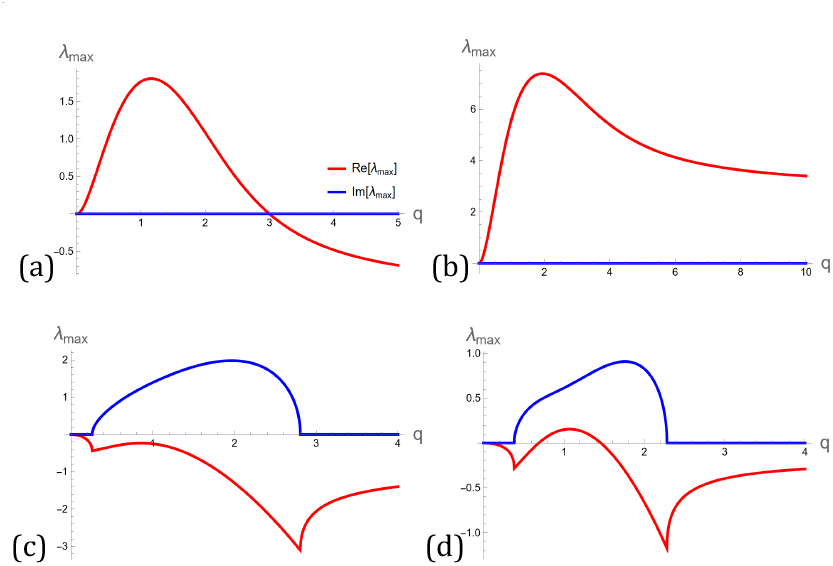
Linear stability dispersion curves, with 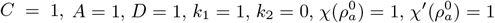, showing (a) segregation instability, (b) elastic contractile instability, (c) stable wave and (d) unstable wave.

If *λ*_*i*_(*q*) are distinct eigenvalues of **M**(*q*) and **v**_*i*_(*q*) are the corresponding (linearly independent) eigenvectors, then we can write the general solution as

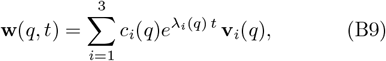

where the coefficients *c*_*i*_(*q*) are the projections of the initial data **w**(*q*, 0) along the respective eigenvectors: (*c*_1_(*q*) *c*_2_ (*q*) *c*_3_ (*q*))^*T*^= **V**(*q*)^*−*1^**w**(*q*, 0), where **V**(*q*) := **[v**_1_(*q*) **v**_2_(*q*) **v**_3_(*q*)] is the matrix containing the eigenvectors as columns.

## Appendix C: Linear Instabilities and Phases

Our first observation is that *λ*_1_ is real and *λ*_1_ *<* 0 for all *q*, since *k*_1_ *>* 0 and *D >* 0 by definition. Hence, all modes corresponding to the first eigenvalue *λ*_1_ are asymptotically stable (i.e., monotonically decaying). Secondly, we observe that *Re*[*λ*_3_] ≥ *Re*[*λ*_2_] for all *q*, i.e., *λ*_*max*_ = *λ*_3_.

Hence, the largest eigenvalue 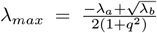 determines the asymptotic stability of the linearized system, i.e., the ‘linear’ phases. We give the definitions of various phases for 1D case in Table I.

**TABLE I.** Definition of the instabilities and phases in the linear analysis.

| (In)stability description | Definition |
| --- | --- |
| Mechanical stability of the elastomer | $\tilde{B} + \mu > 0$ |
| Mechanical instability of the elastomer | $\tilde{B} + \mu \leq 0$ |
| Segregation instability | $\tilde{B} + \mu > 0, \lambda_b > 0, \lambda \geq 0$ |
| Monotonic stability | $\tilde{B} + \mu > 0, \lambda_b > 0, \lambda < 0$ |
| Damped oscillations | $\tilde{B} + \mu > 0, \lambda_b < 0, \lambda_a < 0$ |
| Growing oscillations (unstable) | $\tilde{B} + \mu > 0, \lambda_b < 0, \lambda_a > 0$ |
| Sustained oscillations (swap, travelling waves) | $\tilde{B} + \mu > 0, \lambda_b < 0, \lambda_a = 0$ |

Based on the definitions given in Table I, we can construct various linear stability phase diagrams, which, in principle, depend on the wave vector magnitude *q*. The phase diagrams presented in the main text are constructed assuming *q* to be small, i.e., in the long wavelength limit.

### C1. Segregation

When *λ*_*b*_ *>* 0, then *λ*_2,3_ are real, hence, we have the monotonic stable/unstable phases. The onset of segregation instability is dictated by the sign change of *λ*_*max*_ from negative to positive.

The fastest growing mode *q*_*s*eg_ is where *∂*_*q*_*λ*_*max*_ = 0. With 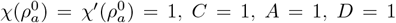 and *k*_1_ = 1, we find

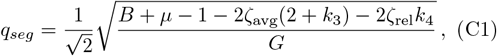

where

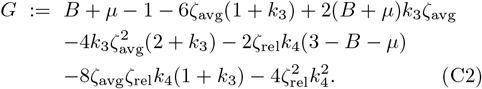

We observe that *λ*_*max*_ = 0 when 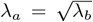, which, using (B7b), implies that

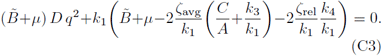

The (positive) solution to this equation

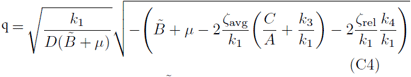

is real and non-zero for 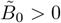 if and only if

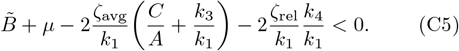

Hence, if the condition (C5) is met, *λ*_3_ becomes non-negative in the long wavelength limit, thus, triggering segregation instability. The characteristic width of the segregated regime is 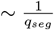.

## Appendix D: Nonlinear Analysis of governing equations

These scaling forms imply the following for the derivatives,

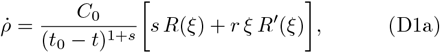

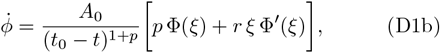

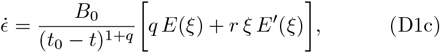

and

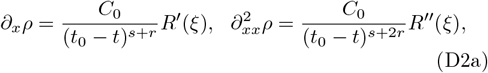

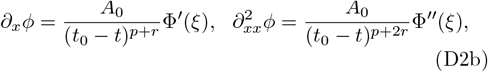

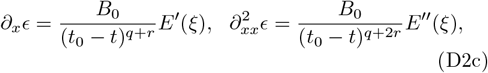

where, (·)^*′*^ denotes differentiation with respect to *ξ*.

We substitute the similarity forms (13), (D1) and (D2) into the system (11), and ignore diffusion as it cannot produce any finite time singularity. The *ϵ*-equation gives,

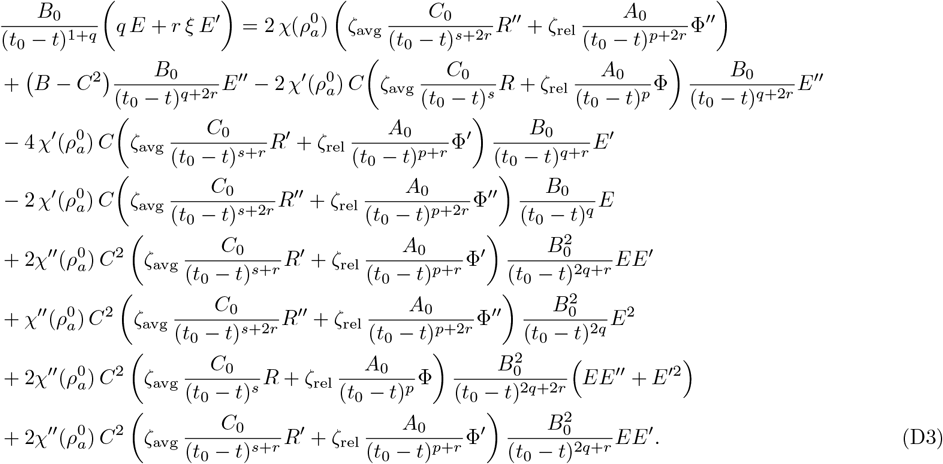

The *ρ*-equation gives,

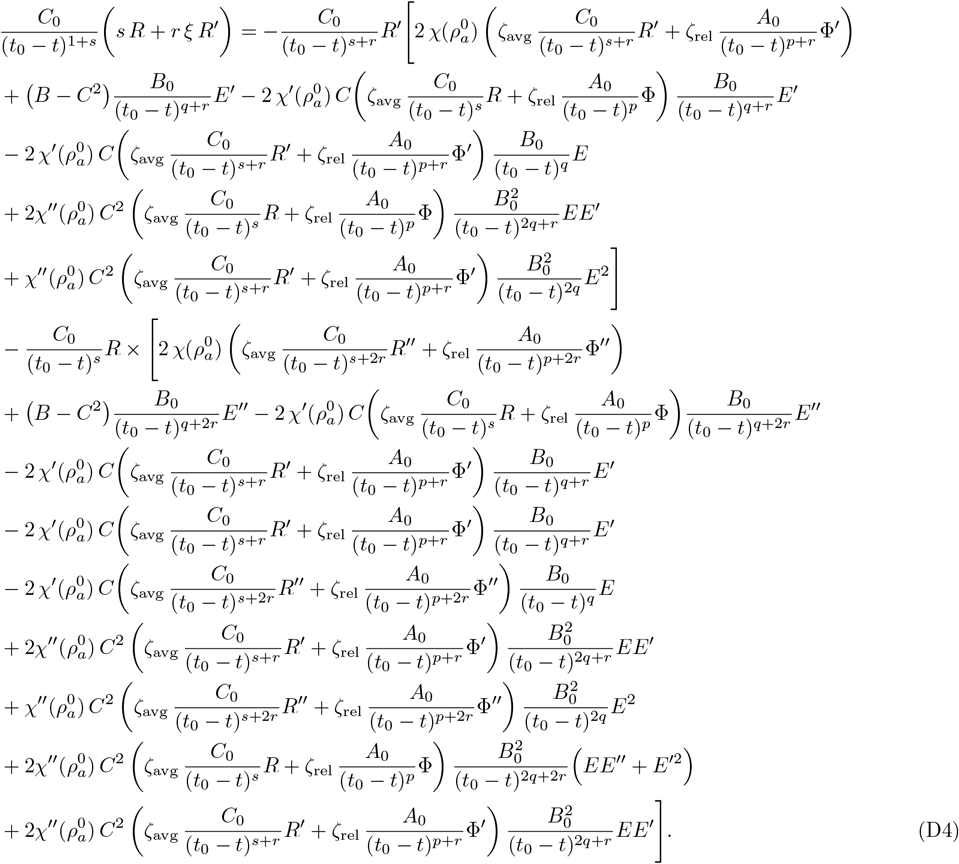

The *ϕ*-equation gives,

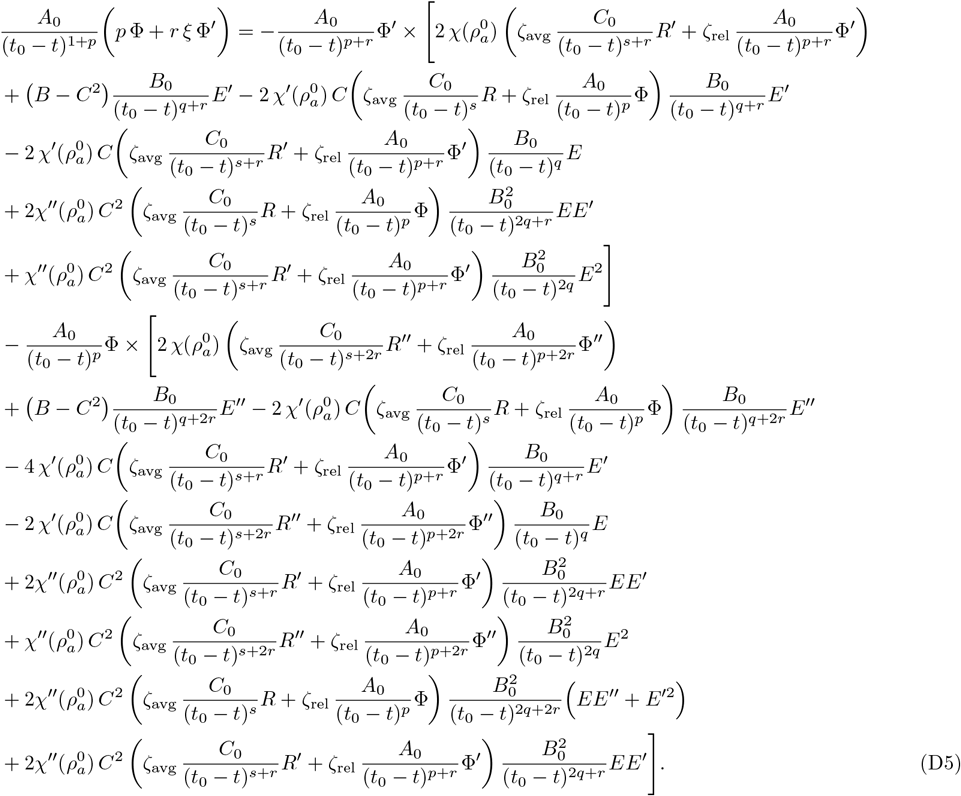

Then by balancing the powers of (*t*_0_ − *t*) on the LHS and *χ*^*′′*^ terms on the RHS, equations (15) in the main text are obtained.

## Appendix E: Kinematics and Balance Laws for 1D Singular Fields

### E1. Compatibility condition

The bulk displacement field *u*(*x, t*) is continuous at the singular point *x* = *x*_0_(*t*), i.e.,

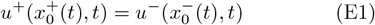

where 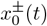 represent the limiting points as one approaches the singularity from the right and the left, respectively, and similarly, *u*^*±*^ represent the corresponding limiting values of the displacement. Taking time derivative of the above equality implies that

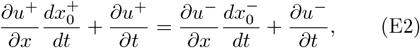

i.e.,

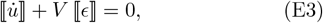

where 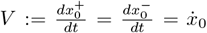 denotes material velocity of the singular point. Thus, the spatial velocity of the singularity is then defined by 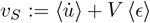.

### E2. Divergence Theorem

Consider a piece-wise smooth bulk scalar field *f* on an arbitrary closed interval [*a, b*] ⊂ ℝ which suffer jump discontinuity ⟦*f*⟧ at a point *x*_0_ ∈ [*a, b*]. Let us partition the interval [*a, b*] into subintervals 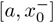 and 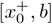 such that 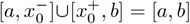 and 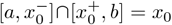.

Hence,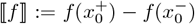. Since *f* is smooth within each of these subintervals, where the standard 1D divergence theorem (a.k.a. fundamental theorem of calculus) applies:

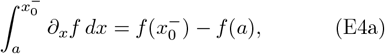

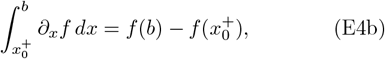

Adding these two identities gives 1D divergence theorem for the piece-wise smooth field *f* over [*a, b*]:

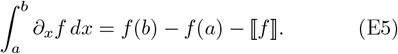

### E3. Transport Theorem

Next, consider a moving singular point *x*_0_(*t*) ∈ [*a, b*] ≡ Ω, with material speed 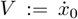. Consider a small *ε*-neighbourhood Ω_ε_ ⊂ Ω of *x*_0_, such that points within Ω_ε_ are parametrized by *x* = *x*_0_ +*ψ*(*t*), with *ψ* = 0 characterizing *x*_0_, and −*ε < ψ*(*t*) *< ε*, ∀*t*, with a constant *ε >* 0. Further, partition Ω_ε_ into subintervals [−*ε, ψ*^*−*^(*t*)] and [*ψ*^+^(*t*), *ε*], such that [−*ε, ψ*^*−*^(*t*)] ∪ [*ψ*^+^(*t*), *ε*] = [−*ε, ε*] and [−*ε, ψ*^*−*^(*t*)] ∩ [*ψ*^+^(*t*), *ε*] = *ψ*(*t*). Hence, 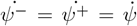, and, moreover, 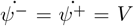 for *ψ* = 0. In this proof, we have closely followed [67].

Then, for a piece-wise smooth bulk scalar field *f* on [*a, b*] which suffer jump discontinuity ⟦*f*⟧ at *x*_0_(*t*) ∈ [*a, b*],

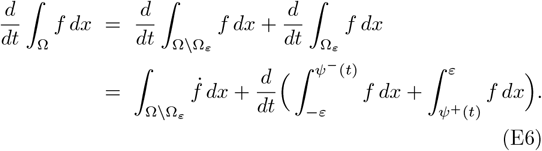

Using the Leibniz integral rule for smooth fields valid on each subintervals, we obtain

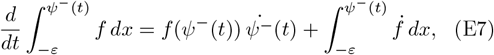

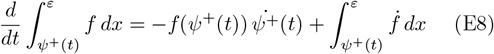

Substituting these two integrals into (E6) and then taking the limit *ε* → 0, imply the following transport theorem for the 1D piece-wise smooth field *f* :

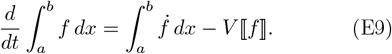

### E4. Mass Balance

Let the bulk and singular mass densities be *ρ* and 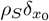, respectively, where 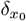 denotes the Diract distribution supported at *x*_0_, the piecewise smooth bulk mass flux be *j* with jump discontinuity ⟦*j*⟧ at *x*_0_, and the bulk and singular mass turnover (due to binding and unbinding) be Π and 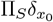, respectively. Then, the equation for the global balance of mass reads

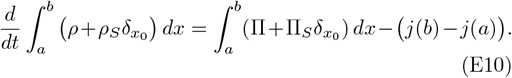

Using the theorems (E5) and (E9), this equation yields

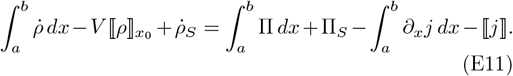

The arbitrariness of the domain [*a, b*] and the point *x*_0_ then yield the local laws

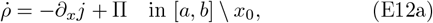

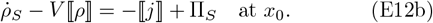

### E5. Force Balance

Let the piecewise smooth bulk (scalar) stress field be *σ*. Then, the global linear momentum balance equation, with environmental viscosity and in absence of inertia, is

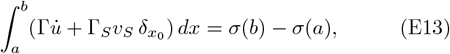

where the left hand side represents the total frictional force. Then, using the theorem (E5), and arbitrariness of the domain [*a, b*] and the point *x*_0_, we obtain the local laws

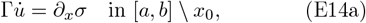

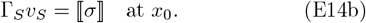

**FIG. E8.**
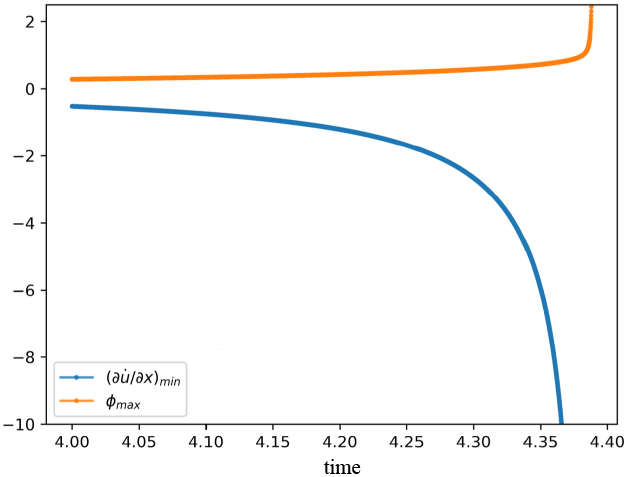
Strain rate 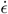 blows up faster than the segregation field *ϕ*, as the singularity is approached.

**FIG. E9.**
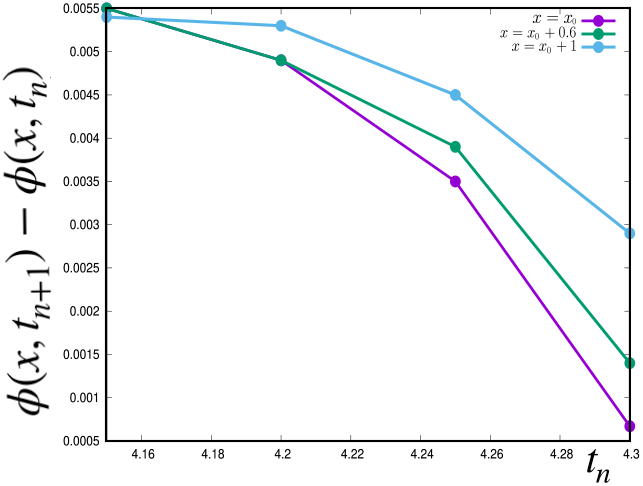
Cauchy convergence of *ϕ*(*x, t*) in time *t* as the singularity is approached; points closer to the blow-up location *x*_0_ diverge faster.

## Appendix F: Movie Captions

### F1. Segregation of a binary mixture of stresslets with density peak co-localization

**[Movie 1](https://drive.google.com/file/d/14CVxT7SFx-76mTdeD2PDLCU-ONjg75jI/view?usp=sharing)**. This movie shows segregation, followed by the formation of singularities in the profiles of *ρ, ϕ* and *ϵ* of the stresslets, where the density peaks co-localize. Initial conditions are homogeneous unstrained state with noise. Both stresslets are of catch bond type. Parameters set at 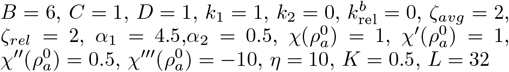.

### F2. Segregation of a binary mixture of stresslets with density peak separation

**[Movie 2](https://drive.google.com/file/d/16a1KWvMt95U8zEvvgcskpv9YLXXqApWs/view)**. This movie shows segregation, followed by the formation of singularities in the profiles of *ρ, ϕ* and *ϵ* of the stresslets, where the density peaks separate. Initial conditions are homogeneous unstrained state with noise. One stresslet is of a catch bond type and the other is of a slip bond type. Parameters are set at 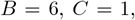 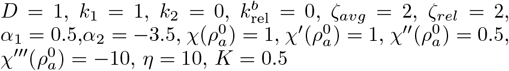.

### F3. Dynamics of singularity formation

**[Movie 3](https://drive.google.com/file/d/1nAkaJR_G6udob4C0n7vA6DdXK_b3O7D4/view)**. This movie (left panel) shows the approach to a finite time elastic singularity, and (right panel) shows its physical resolution by a steric term (*B*_3_ ≠ 0). Parameters are set at 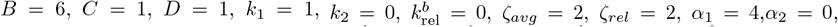 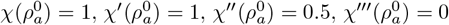 (for left panel), 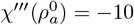 (for right panel), *η* = 10, *K* = 0.1, *L* = 10.

### F4. Coarsening Dynamics: catch bond with turnover

**[Movie 4](https://drive.google.com/file/d/1ci_D42aP20kXC45w0JwtzV5Nisizz1hS/view)**. This movie shows the coarsening dynamics of elastic singularities, in a system where both stresslets have catch bonds. At first, the singularities merge quickly following which the coarsening slows down, showing signs of dynamical arrest. Parameters are set at 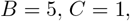 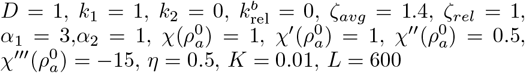.

### F5. Coarsening Dynamics: catch bond without turnover

**[Movie 5](https://drive.google.com/file/d/1XYkDBGoiNu1z5DrmqNfOgcsHBNfQXiD2/view)**. The configurations generated in the previous numerical simulation of coarsening dynamics of the species with catch bonds at *t* = 100, is further evolved without turnover. The singularities broaden and merge faster in the absence of turnover and even undergo splitting on occasion. Parameters are set at 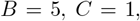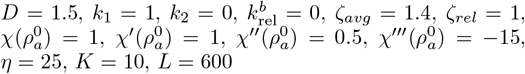.

### F6. Coarsening Dynamics: slip bond with turnover

**[Movie 6](https://drive.google.com/file/d/1XYkDBGoiNu1z5DrmqNfOgcsHBNfQXiD2/view?usp=sharing)**. This movie shows the coarsening dynamics of the singularities where one stresslet has a catch bond and the other, a slip bond. Parameters are set at 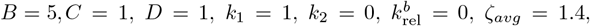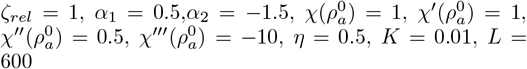.

The pseudospectral code and the analysis notebook used to create the Movies 1-6 can be accessed **[here](https://github.com/saptarshid999/active_force_patterning)**.

## Notes

https://arxiv.org/abs/2409.09050

### Competing Interest Statement

The authors have declared no competing interest.

### Summary of Updates

Revised Abstract, Introduction, Discussion, few sections of the paper removed.

